# Directional Sensing by Eukaryotic Receptors

**DOI:** 10.1101/2024.11.22.624862

**Authors:** Andrew Goetz, Purushottam D. Dixit

## Abstract

*Directional sensing* enables eukaryotic cells to detect spatial gradients of extracellular ligands by comparing local signaling activity to an internal set point. While this capability is typically attributed to signaling pathways downstream of cell surface receptors, we present a reaction–diffusion model to argue that receptors themselves can perform directional sensing via a mechanism we term Localized Activity and Global Sensitization (LAGS). LAGS integrates common receptor signaling motifs: lateral diffusion, basal activity, and active receptor degradation. Without diffusion, receptor activity locally adapts to a ligand-independent set point via degradation-mediated integral feedback, leading to receptor depletion in ligand-rich regions. The introduction of lateral diffusion redistributes primarily ligand-free receptors across the cell surface, creating a spatial mismatch between local activity and feedback. This mismatch drives polarized receptor activity relative to the set point, while rapid ligand equilibration and receptor degradation protects it from diffusion. The model predicts an optimal diffusion coefficient that maximizes receptor polarization, and shows that cells respond to relative – not absolute – changes in ligand levels. Cooperative receptor interactions further amplify the polarized response. Finally, stochastic analysis identifies an optimal set point that maximizes the signal-to-noise ratio in polarization. A survey of kinetic parameters across diverse receptor systems supports LAGS as a general mechanism for receptor-mediated directional sensing.

**Significance statement:** Cells navigate their spatially heterogeneous environments by detecting spatial gradients of external signals – a process known as *directional sensing*. While traditionally attributed to downstream signaling pathways, we argue that cell surface receptors can themselves perform directional sensing by leveraging lateral diffusion, basal activity, and active receptor degradation. Using a reaction–diffusion model, we identify a general mechanism of directional sensing which we call Localized Activity and Global Sensitization (LAGS). In LAGS, degradation-driven integral feedback and receptor diffusion together generate robust, spatially polarized receptor activity. Our results suggest that directional sensing can emerge directly from receptor-level dynamics, revealing a broadly applicable strategy encoded in the biophysics of diverse signaling systems.

## I. INTRODUCTION

Eukaryotic cells display directionally asymmetric behaviors such as chemotaxis and chemotropism (1–11). These behaviors often begin with *directional sensing*, which is the cells’ ability to determine their orientation within spatial gradients of extracellular ligands (2–6, 12–15).

A salient feature of directional sensing circuits in eukaryotic cells (6, 16–18) is the establishment of a ligand-independent set point of signaling activity (18–20). In the presence of spatially uniform ligand stimulation, signaling activity is independent of the external ligand concentration. In contrast, when gradients are imposed, regions of the cell exposed to above-average ligand levels exhibit signaling above this set point, while those sensing below-average ligand levels show reduced activity. These deviations encode positional information that initiates asymmetric cellular responses.

Prevailing theoretical models of eukaryotic directional sensing attribute these computations to signaling networks downstream of ligand-sensing receptors, such as the RAS/RAF cascades or lipid signaling (2, 3, 13, 21–26). In contrast, the contribution of receptor-level processes such as diffusion, basal activation, and degradation remain less explored (27–30).

Yet, several lines of evidence suggest receptors actively participate in directional sensing. In spatial gradients of epidermal growth factor (EGF), EGF receptors (EGFRs) are preferentially internalized at the leading edge of migrating cells, despite their uniform distribution on the plasma membrane (31, 32), underscoring the role of lateral diffusion. Reducing receptor diffusion, for example, by depleting membrane cholesterol (33, 34), impairs gradient sensing (35–37) even when receptor signaling and trafficking remain unaffected (36). Similarly, blocking endocytosis or receptor degradation disrupts chemotaxis (32, 38–42), and mutations that abrogate active receptor degradation reduce chemotactic accuracy despite increasing total receptor abundance (41) (which is otherwise expected to lower sensing noise (27)). Finally, in many cases, adaptation in downstream signaling components (e.g., Akt, Erk, or lipid mediators) reflects upstream adaptation in active receptor levels (43–45). These findings suggest that receptor-level processes may be integral to directional sensing. Although such processes have been proposed to enhance sensing in uniform ligand fields (46, 47), their role in spatial sensing remains largely unexplored.

To investigate how receptors contribute to spatial gradient sensing, we develop a simple reaction–diffusion model that incorporates three features common to many receptor systems: lateral diffusion in the membrane (48–50), ligand-independent basal activity (51, 52), and active receptor degradation (53–56). Using this model, we uncover a receptor-level mechanism for directional sensing, which we term **L**ocalized **A**ctivity and **G**lobal **S**ensitization (LAGS).

When receptor diffusion is slow, degradation of active receptors implements an integral feedback loop (19, 20, 44), driving local receptor activity toward a ligand-independent set point. This leads to receptor depletion in ligand-rich regions, but no spatial variation in activity, preventing directional sensing. Introducing lateral diffusion causes redistribution of predominantly ligand-free receptors toward depleted regions, disrupting the local balance between activity and feedback. The resulting mismatch generates polarized receptor activity - above the set point in high-ligand regions and below it in low-ligand regions - despite the system remaining adapted globally.

The model predicts that active receptor polarization encodes relative rather than absolute ligand concentrations (44, 57), and operates most effectively at subsaturating ligand levels - consistent with physiological ranges (58). Cooperative ligand binding enhances polarization, allowing active receptors to amplify shallow input gradients. A stochastic analysis of a coarse-grained model identifies an optimal network set point that maximizes the signal-to-noise ratio in receptor polarization.

Altogether, our model reveals how directional sensing may arise at the receptor-level. It explains several previous observations in cell migration and polarity and suggests new, testable predictions. A survey of kinetic parameters across receptor families supports LAGS as a broadly applicable mechanism for directional sensing.

## II. RESULTS

### A. A reaction/diffusion model for directional sensing by receptors

To understand the combined role of receptor-level processes on directional sensing, we develop a one-dimensional reaction-diffusion model of ligand-mediated signaling that captures key features of receptor signaling pathways such as RTKs and GPCRs. Although a simplification of the complex three-dimensional cell, a one-dimensional model is sufficient to illustrate the asymmetric distribution of receptors on the cell surface in response to spatial ligand gradients. Indeed, several previous works have used one-dimensional models (2, 3, 22) to study directional sensing.

In our model, we imagine a cell that exists between *X* = 0 and *X* = *l*_*C*_ where *l*_*C*_ is the length of the cell. The schematic of the model and the reactions that can happen on the cell surface are shown in Figure 1. Briefly, *R*(*X*) and *A*(*X*) denote the abundance of inactive and active receptors at position *X*. The receptors diffuse within the bounds of the cell with a diffusion constants *D* (48–50). We denote the externally imposed gradient of the ligand as *L*(*X*). For simplicity, we assume that the gradient is not perturbed via interactions with the cell. Inactive receptors are uniformly delivered to the cell surface at a constant rate *k*_delivery_. Receptors are activated at a ligand-independent basal rate *k*_*ϕ*_ (51, 52) and a ligand-induced rate *k*_1_. Active receptors lose activity at a rate *k*_−1_. Finally, we assume that active receptors are internalized and degraded at a rate 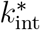 (53–56).

**FIG. 1.**
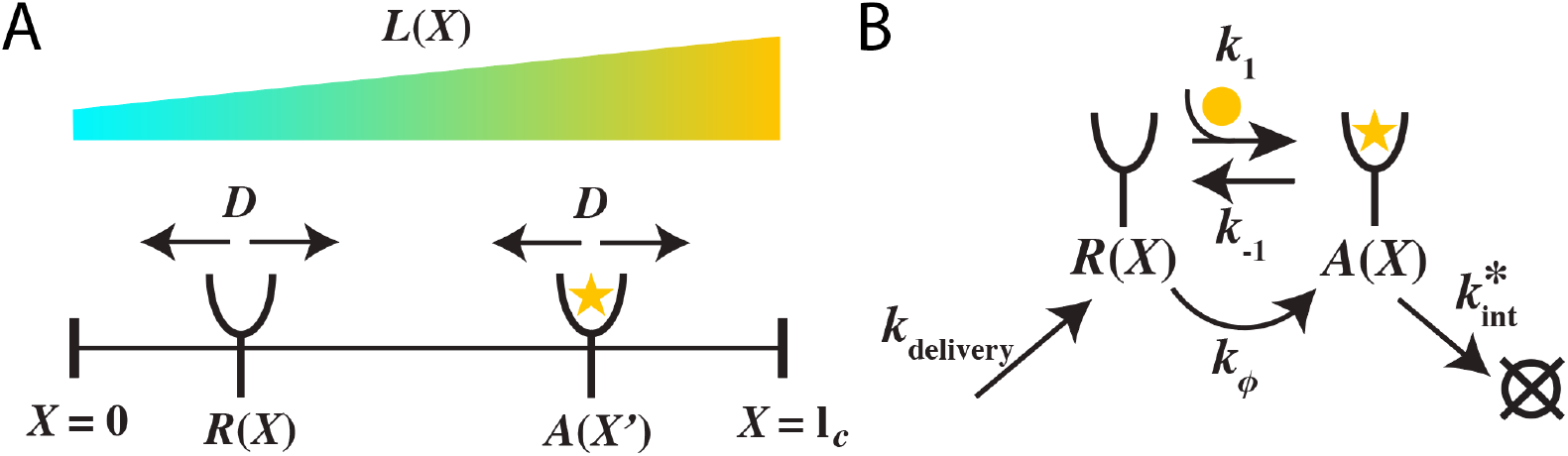
Schematic of the reaction-diffusion model. (A) Inactive and active receptors diffuse along the cell surface between coordinates *X* = 0 and *X* = *l*_*C*_ with diffusion constant *D*. An external gradient of ligand concentration *L*(*X*) is imposed. (B) Reactions that take place at every location *X* in the model.

The dimensionless reaction/diffusion system is given by (see Supplementary Materials Section I):

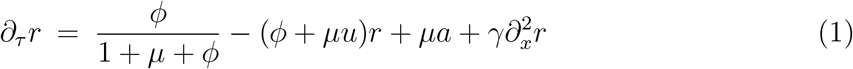

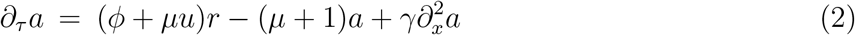

where 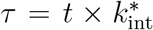 ensionless time and *x* = *X/l*_*C*_ is the dimensionless position. 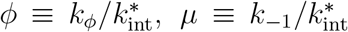, and 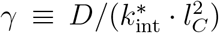are dimensionless kinetic parameters. We have defined dimensionless ligand concentration *u*(*x*) = *L*(*x*)(*k*_1_*/k*_−1_). We have also introduced *r* = *R/R*_0_ and *a* = *A/R*_0_ as dimensionless abundances where *R*_0_ is the receptor abundance in the absence of ligand exposure.

#### Parameter ranges

To investigate how active receptor polarization depends on network parameters, we first identified realistic ranges for the dimensionless kinetic rates. The length of a mammalian cell is typically *l*_*C*_ *≈* 10 − 60*µm* (59), the lateral diffusion constants of receptors range between *D ≈* 0.015 − 0.25*µm*^2^*s*^−1^ (48–50), the internalization rate of active receptors is 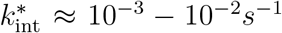 (54, 56). Ligand dissociation rates are typically *k*_−1_ *≈* 10^−3.5^ − 10^−0.5^*s*^−1^ (44, 60). In our analysis below, we use dimensionless parameter ranges that are generously inclusive of these parameters. Specifically, we choose *µ ∼* 0.1 − 1000 and *γ ∼* 0.001 − 1. Based on basal phosphorylation levels for realistic receptor counts (51, 61), we estimate *ϕ ∼* 0.01 − 10.

In what follows, we assume that the right half of the cell is exposed to a higher ligand concentration compared to the left half. Specifically, we impose a small linear ligand gradient across the cell surface, represented by *u*(*x*) = *u*_0_(1+*gx*) where *g >* 0 is the gradient and *u*_0_ is the background ligand concentration. To streamline the discussion, unless stated otherwise, we use *µ* = 100, *γ* = 0.015, and *ϕ* = 0.1.

Using our model, we study the spatial distribution of active receptors at steady state. Directional sensing in our model can be understood as follows. As shown in Figure 2B, when diffusion is negligible (*γ ∼* 0), active receptor levels at all locations on the cell adapt to a ligand-independent set point via a degradation driven integral feedback (Figure 2A) (18– 20, 44). The implementation of the integral feedback can be seen by observing that in the absence of diffusion, the total receptor abundance

**FIG. 2.**
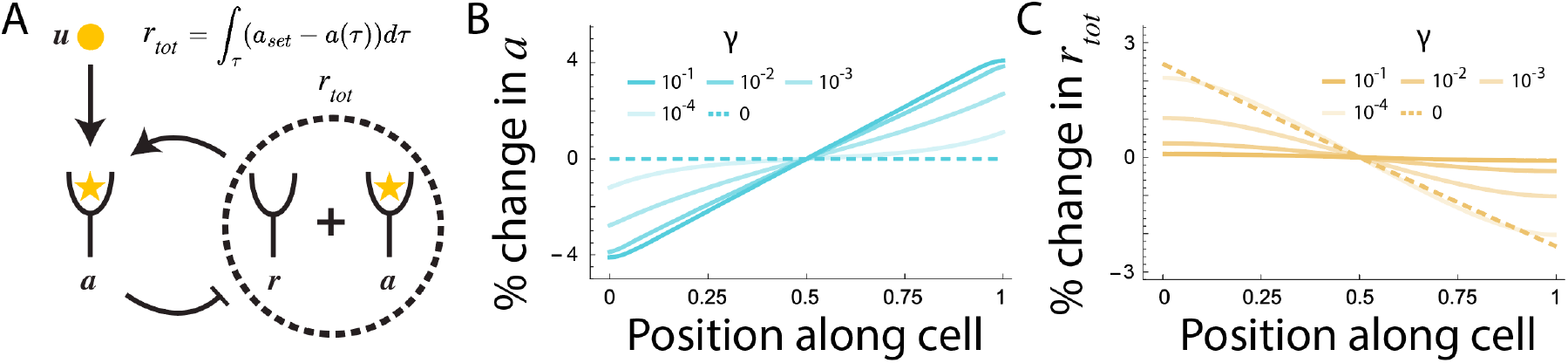
Integral feedback and spatial receptor profiles. (A) Schematic of the integral feedback in the model. (B and C) The % change in active receptor count (B) and the total receptor count (C) plotted as a function of position along the cell surface for different values of the diffusion constant. *r*_tot_ is normalized with respect to its value the center of the cell and *a* is normalized with respect to the network set point *a*_set_. In both cases, we have used a gradient of *g* = 10%.

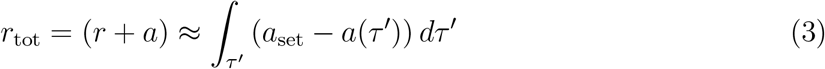

at every location on the cell surface is the local *integrator* species that achieves adaptation in the active receptor levels (18–20, 44). Here, we recognize

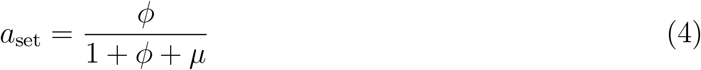

as the ligand-independent set point of the network.

Figure 2C shows that as a result of receptor degradation, the total receptor abundance in regions of the cell exposed to high extracellular ligand levels is lower compared to regions of the cell exposed to lower ligand levels. Introduction of receptor diffusion imbalances the feedback. When *γ >* 0, the gradient in total receptor count is dissipated via a non-equilibrium flow of receptors from regions of low ligand exposure to regions of high ligand exposure. As a result, regions exposed to high ligand levels reach a steady state receptor activity above the network set point while regions experiencing low ligand levels reach an activity below the set point (Figure 2B). This way, different regions of the cell perform directional sensing using local receptor activity. Specifically, regions with receptor activity above *a*_set_ are closer to the leading edge and vice versa for regions with activity below *a*_set_. Notably, this computation can be performed independent of the background ligand concentration *u*_0_, a hallmark of directional sensing.

### B. Diffusion, dissociation, and degradation are key components of LAGS

In order to compare directional sensing across different sets of parameters, we quantify active receptor polarization (ARP) as the difference in active receptor abundance between the two ends of the cell normalized by the network set point:

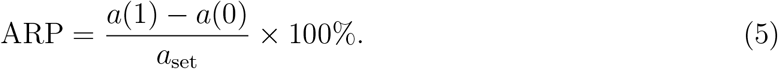

The normalization with respect to *a*_set_ allows us to compare different parameter sets that may lead to different set points and therefore different overall response magnitudes. In addition to network parameters, ARP also depends on the background ligand levels (see Figure 4 below). Unless stated otherwise, we report the ARP at the optimal background ligand level *u*_0_ that corresponds to its maximum (Max ARP).

Figure 3A shows the ARP as a function of diffusion constant *γ* and dissociation rate *µ*. For fixed *γ*, ARP sharpens with an increase in *µ*. This is because rapid ligand equilibration protects the local receptor activity from diffusing along the cell surface. At intermediate diffusion rates (*γ ∼* 0.01 − 0.1), ARP is robust to decrease in *µ* when *µ ∼* 1. In this regime, active receptor profile is localized predominantly due to receptor degradation. This way, a combination of rapid equilibration and receptor degradation localizes activity (LA).

**FIG. 3.**
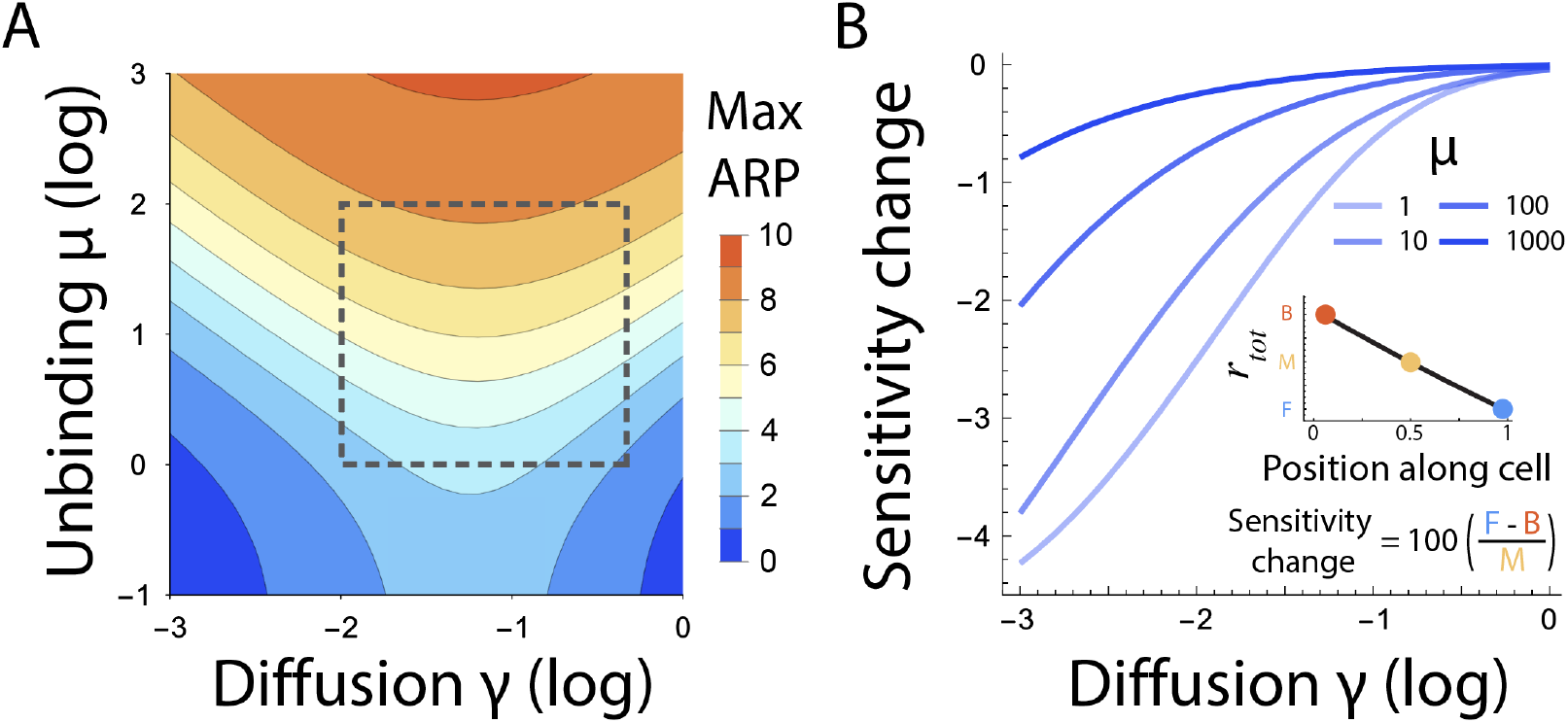
Mechanism of LAGS. (A) Active receptor polarization as a function of diffusion constant *γ* and dissociation rate *µ*. Dashed squares indicate typical parameter ranges for RTKs and GPCRs based on literature derived kinetic parameters. (B) Change in sensitivity (total receptor count) at the background ligand level corresponding to the max ARP as a function of *γ* for different values of *µ*. See inset for the definition of sensitivity change. A gradient of *g* = 10% is used in both plots.

Notably, ARP has a non-monotonic dependence on diffusion. The decrease in ARP at high diffusion rates (*γ >* 0.1) is due to diffusion-driven smearing of receptor activity. In contrast, a counter intuitive increase in ARP with diffusion at low diffusion values can be understood in terms of sensitization. Figure 3B shows that diffusion facilitates redistribution of receptors along the cell surface, leading to global sensitization (GS) towards the ligand. A more uniform receptor distribution creates a higher mismatch between degradation-based feedback and activity, leading to a more sharply polarized activity.

Figure 3A shows typical non-dimensional parameter ranges for RTKs and GPCRs, based on a literature survey (dashed black lines) (44, 48–51, 54, 56, 60, 61). These ranges suggest that the kinetic parameters of chemosensing networks may be tuned to exploit an optimal diffusion constant for receptor polarization.

### C. Relative sensing and response amplification

Signaling networks with integral feedback often exhibit relative sensing in temporal ligand gradients (44, 62). To test whether this principle extends to spatial gradients, we analyzed active receptor polarization (ARP) across varying gradient strengths *g* and background ligand levels *u*_0_ (Figure 4A). We find that ARP is largely insensitive to background ligand levels *u*_0_, but responds robustly to the relative gradient *g*, a hallmark of relative sensing (44, 57). Notably, for realistic parameters, the background concentration that elicits maximal ARP (*u*_0_ *∼* 10^−2^) is two orders of magnitude below the equilibrium dissociation constant of the ligand. This concentration range is also significantly lower than those typically used in *in vitro* experiments (see for example, (63–65)) but matches ligand concentrations observed *in vivo* (58). These results suggest that receptors themselves can perform relative sensing in spatial gradients, potentially shaping downstream responses and cellular behavior such as chemotaxis. Moreover, optimal relative sensing occurs in subsaturating ligand concentrations.

**FIG. 4.**
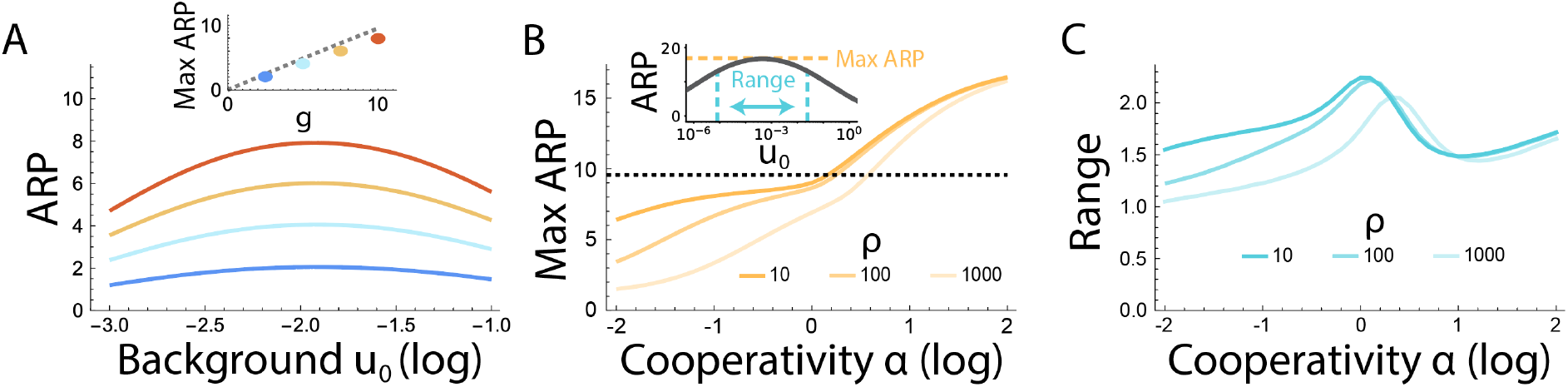
Relative sensing and amplification. (A) ARP as a function of background ligand concentration *u*_0_ for different values of gradients: *g* = 2.5%, 5%, 7.5%, and 10%. Inset shows the maximum ARP as a function of *g*. (B) Maximum ARP plotted as a function of cooperativity *α* for different values of dimensionless dimerization rate *ρ*. (C) The range of relative sensing (see inset in B) as a function of cooperativity *α* in receptor dimerization plotted for different values of dimensionless receptor dimerization rate *ρ*.

The inset of Figure 4A shows that the ARP depends linearly on the imposed gradient 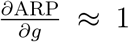. Can polarized receptor activity achieve response amplification 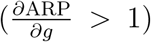? A key feature of many receptor-ligand systems is receptor oligomerization following ligand binding (66). Moreover, oligomerization can sharpen or flatten activity response depending on cooperativity in receptor-receptor interactions (67, 68). To understand how receptor oligomerization may affect ARP, we introduce receptor dimerization in our model. We consider two types of receptor dimers *d*_1_ = *r · a* and *d*_2_ = *a · a*. The total activity is defined as *A* = *d*_1_ + 2*d*_2_. This modification introduces three new dimensionless constants: the dimerization rate *ρ*, the undimerization rate *ω*, and the cooperativity *α* (see Supplementary Materials Section II). *α >* 1 signifies positive cooperativity as it favors the formation of the *a · a* dimer over the *r · a* dimer and vice versa for *α <* 1. Figure 4B shows that positive cooperativity in receptor oligomerization can lead to receptor activity gradient that is amplified (up to *∼* 60%) compared to the imposed gradient. Notably, the gain of ARP was constant across a broad range of imposed gradients *g* (see Supplementary Figure S1). Higher order clusters (not examined here) are likely to produce even sharper response amplification (69). In contrast, negative cooperativity resulted in a response with a gain *<* 1. Notably, Figure 4C shows that the amplification comes with a tradeoff; the range of relative sensing (background ligand concentration range over which the ARP is greater than 80% of its maximum, inset of Figure 4B) shrinks with both positive and negative cooperativity.

### D. Stochastic modeling identifies an optimal network set point

Biochemical networks are often subject to stochastic fluctuations due to small molecule numbers. To understand how stochastic fluctuations may affect directional sensing, we employ a simplified two compartment model (Figure 5A, see Supplementary Materials Section III). A recent work used a similar analysis to characterize stochasticity in models of directional sensing in yeast mating (70). The model characterizes the full joint distribution of all chemical species (active and inactive receptor abundance in the two compartments) as a function of kinetic parameters. Surprisingly, this simplified analytical model qualitatively reproduces all predictions of the full reaction/diffusion model (see Supplementary Materials Section III). For the stochastic model, the signal-to-noise ratio (SNR) in receptor polarization is given by (see Supplementary Materials Section III):

**FIG. 5.**
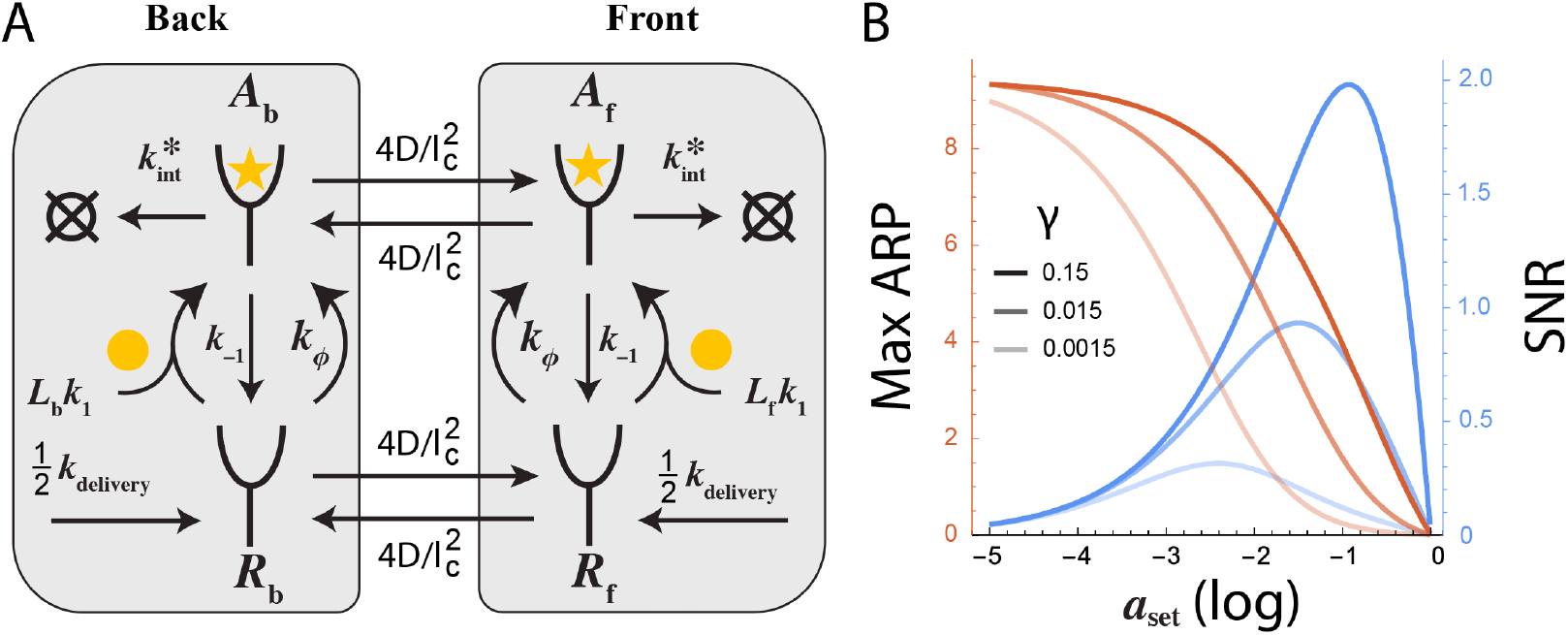
Stochastic analysis identifies an optimal set point. (A) Simplified two compartment model used to investigate stochastic effects. The factor of 4 acting on the diffusion terms result from the discretization process as described in Bernstein (71). (B) Active receptor polarization computed using the deterministic model and signal-to-noise ratio computed using the stochastic model plotted as a function of the basal activity *a*_set_ for different values of the diffusion constant *γ*. We have fixed the receptor count *R*_0_ = 10^5^ receptors per cell.

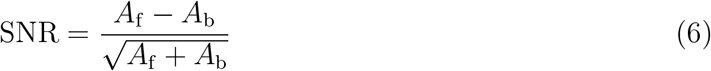

where *A*_f_ and *A*_b_ are absolute receptor activities. In this model, the SNR is proportional to the deterministic ARP with the proportionality constant equal to 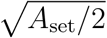 where *A*_set_ = (*A*_*f*_ + *A*_*b*_)*/*2 = *R*_0_*a*_set_*/*2 is the absolute value of the set point (see Supplementary Materials Section III). Consequently, the dependence of the SNR mirrors the dependence of the ARP for most model parameters. Notably, the stochastic model identifies an optimal network set point that is not apparent in the deterministic analysis. In Figure 5B, we plot the SNR (blue) as a function of the set point at *R*_0_ = 10^5^ receptors per cell (61) along with the deterministic ARP (red). The deterministic model predicts that ARP improves monotonically with lowering the network set point. In contrast, the stochastic model shows that enhanced stochastic fluctuations associated with a low set point decreases the signal-to-noise ratio, resulting in optimal performance (in terms of SNR) at intermediate values of the set point.

## III. DISCUSSION

Eukaryotic cells detect and respond to spatial gradients of extracellular signals through a process known as *directional sensing* (2–6, 12–15). Previous theoretical work on directional sensing (2, 3, 13, 21–25) focused on computations downstream of ligand-bound receptors. Here, we demonstrated that cell surface receptors themselves can perform directional sensing by leveraging lateral receptor diffusion, basal activation, and active receptor degradation.

A key feature of our model is the integral feedback implemented through degradation of active receptors (19, 44). Perfect adaptation in this framework assumes that inactive receptors are spared from degradation. While experimental data suggest that inactive receptors do degrade, they typically do so at substantially lower rates than their active counterparts (54, 72). We find that the LAGS mechanism remains robust across a wide range of such *leakiness* in the feedback loop and only fails when degradation rates of active and inactive receptors become comparable (Supplementary Materials Section IV).

How does LAGS relate to prior models of directional sensing in eukaryotic cells? Realistic biological adaptation circuits are inherently leaky, for example, due to factors such as energy dissipation (73, 74) or cell growth and dilution (75). To compensate for leakiness in individual circuits, cells may stitch together multiple adapting modules. For instance, downstream signaling networks often implement local excitation and global inhibition (LEGI) mechanisms that act on activated receptor signals (2, 24). Our results suggest a synergy between LAGS and LEGI: receptor-level adaptation via LAGS can pre-process spatial ligand information before it is further refined by downstream LEGI-type circuits, enabling robust directional sensing through distributed, layered adaptation.

Our model makes several specific and experimentally testable predictions. First, it predicts that reducing receptor diffusion diminishes active receptor polarization (Figure 3). This can be achieved by depleting membrane cholesterol, which slows diffusion without necessarily affecting receptor signaling or trafficking (33, 34, 36). In line with this prediction, cholesterol depletion has been shown to impair chemotaxis across multiple signaling pathways (35–37). Second, the model predicts that weakening the bias toward degradation of active receptors (i.e. increasing the *leak* in the integral feedback) degrades directional sensing at the receptor level (Supplementary Figure S4). Although reducing degradation can increase overall receptor abundance (41) (and potentially reduce polarization noise (27)) such manipulations have been shown to impair chemotaxis (32, 38–42). Third, consistent with imaging data (31, 32), our model predicts uniform receptor abundance along the cell membrane but a polarized pattern of endocytosis biased toward the leading edge (Figure 2, note the extent of endocytosis along the cell is proportional to *a*(*x*)). Finally, the model predicts that cells respond to relative, not absolute, ligand gradients, and that this computation occurs at the level of active receptor polarization. This receptor-level relative sensing may propagate downstream to control cellular behaviors such as the chemotactic index. Notably, relative sensing is most prominent at background ligand concentrations *u*_0_ *∼* 10^−2^, far below the equilibrium dissociation constant and typical levels used in *in vitro* experiments (63–65), but consistent with *in vivo* estimates of ligand levels (58). While previous work demonstrated temporal relative sensing via ligand-induced receptor degradation (44), spatial relative sensing by cell surface receptors remains experimentally untested. Testing these predictions will be a key focus of future work.

Therefore, we believe that LAGS may be a general mechanism employed by cells in their quest to navigate their spatially inhomogeneous environment.

## Supplementary Materials

All model calculations are performed in Mathematica. The scripts are included with the manuscript.

### 1. NON-DIMENSIONALIZATION OF THE MODEL

To derive the non-dimensional equations in the main text, we begin with the reaction diffusion equations describing the monomeric receptor system:

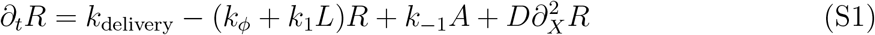

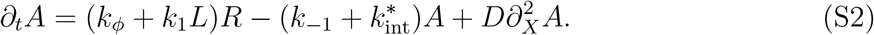

In order to non-dimensionalize these equations, we divide both sides of equations S1 and S2 by 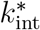, and make use of the reparameterization: 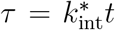, *x* = *X/l*_c_ (*l*_c_ is the length of the cell), *u* = *Lk*_1_*/k*_−1_, *r* = *R/R*_0_ (*R*_0_ is the basal total receptor level, see below), and *a* = *A/R*_0_. From this we obtain:

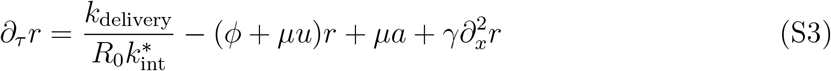

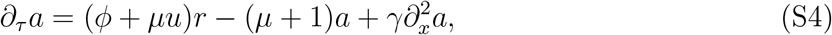

where we identify dimensionless rate constants 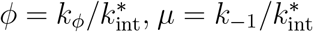, and 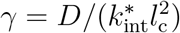. Next, in order to find *R*_0_, we solve equations S3 and S4 at steady state (*∂*_*τ*_ *r* = 0 and *∂*_*τ*_ *a* = 0) with no ligand input (*u* = 0). Here we note the absence of a ligand gradient removes any spatial variation in receptors along the cell, making 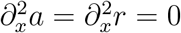. This allows the system to be solved algebraically, giving:

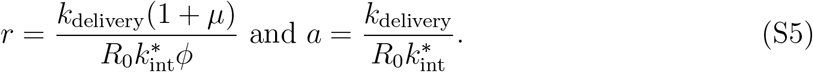

Under this dose-less steady state condition, *R*_0_ = *R* + *A* = *R*_0_(*a* + *r*), so:

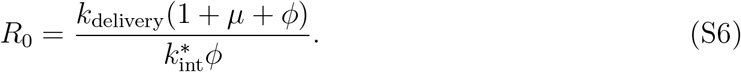

Substituting this expression for *R*_0_ back into equations S3 and S4 gives the following dimensionless equations for the system:

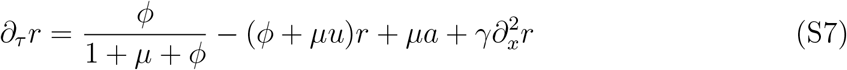

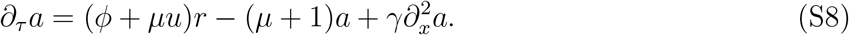

From these equations, we obtain the network set point:

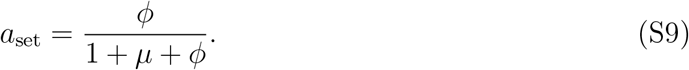

### II. INTRODUCING RECEPTOR OLIGOMERIZATION

The model of receptor dimerization used in this paper is described by the following reaction diffusion equations:

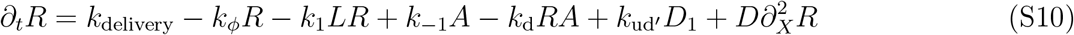

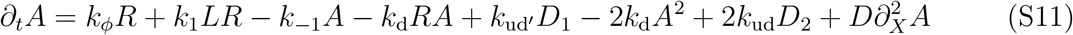

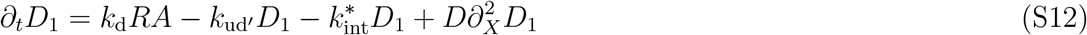

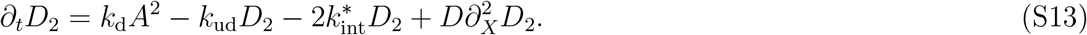

Here, *R* are inactive receptors, *A* are pre-dimers (receptors that are prone to dimerization) that are formed either via basal activation or via ligand binding. *D*_1_ = *R· A* and *D*_2_ = *A· A* are two types of dimers.

In order to non-dimensionalize this model, we divide both sides of equations S10 - S13 by 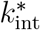, and make use of the re-parameterizations: 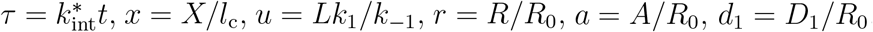, and *d*_2_ = *D*_2_*/R*_0_. From this we obtain:

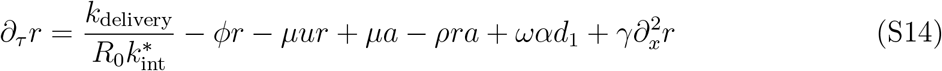

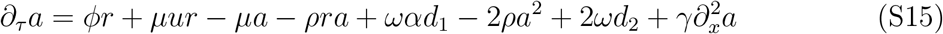

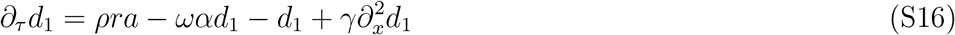

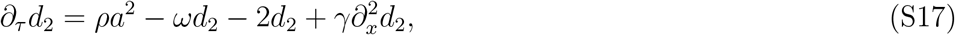

where we define dimensionless kinetic parameters 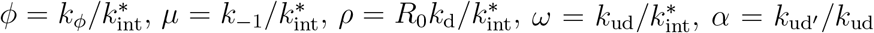, and 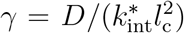. Next we find the steady state solution to the system in the absence of input and obtain *R*_0_ (*R*_0_ = *R* + *A* + *D*_1_ + *D*_2_). Unlike the monomer case, *R*_0_ lacks a tidy analytical expression, so the dimensionless term given by 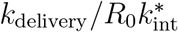 is numerically found for each system prior to solving the reaction diffusion system.

Our model shows that dimerization provides a means for the chemosensing circuit to amplify the ARP beyond the dynamic range of the imposed gradient. While the main text showed this holds true for a fixed imposed gradient of *g* = 10%, the amplification gain from dimerization is in fact largely independent of the imposed gradient (Figure S1).

**FIG. S1.**
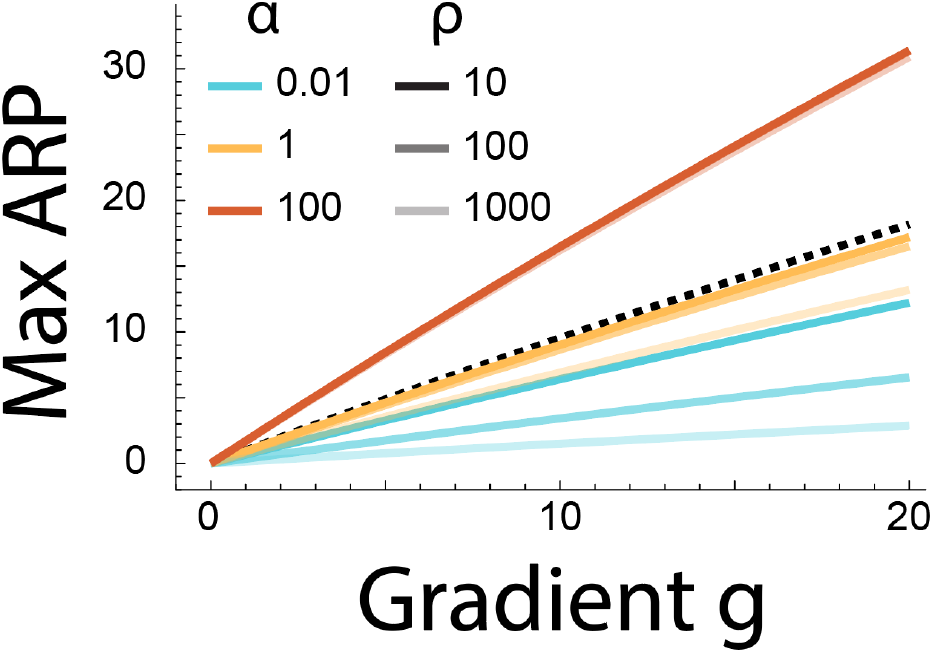
Impact of imposed gradient on gain. Effects of imposed gradient on ARP for different values of dimer cooperativity (*α*) and dimer binding rate (*ρ*). The dashed black line shows the optimal response without amplification.

### III. SIMPLIFIED TWO COMPARTMENT MODEL EXPLORES THE ROLE OF STOCHASTICITY

In our two compartment stochastic model, diffusion is incorporated as a first order reaction which traffics species from one compartment to another. The conversion of the diffusion rate to a first order rate constant in a discretized reaction scheme was described by Bernstein (71): 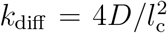where D is the one-dimensional diffusion coefficient and *l*_c_ is the length of the cell. Treating diffusion this way lets us write the two compartment model as a system of ODEs:

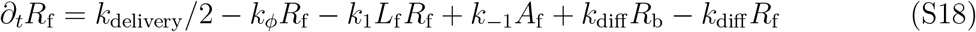

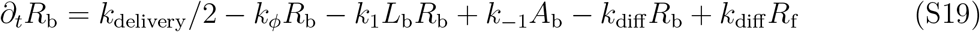

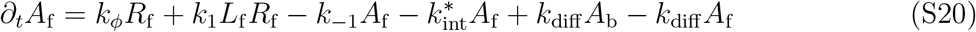

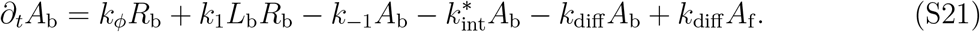

Note that we have scaled the *k*_delivery_ rate in both cell halves to ensure *k*_delivery_ is the total rate of receptor delivery along the entire cell.

Next, in order to non-dimensionalize the model, as above, we divide both sides of the above equations by 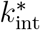, and make use of the re-parameterization, 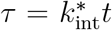. From this we obtain:

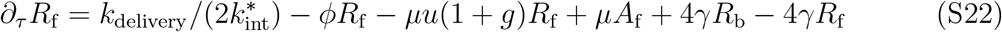

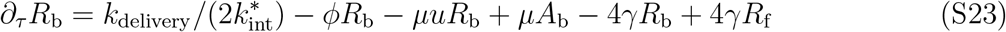

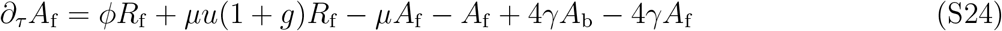

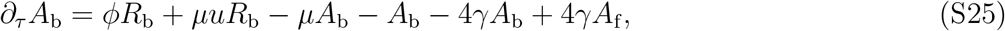

where 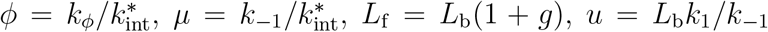, and 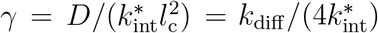.

As above, we define *R*_0_ as the steady state basal receptor count in the absence of the ligand. In order to find *R*_0_, we solve equations S22-S25 at steady state (*∂*_*τ*_ *r* = 0 and *∂*_*τ*_ *a* = 0) with no ligand input (*u* = 0) for *R*_f_, *R*_b_, *A*_f_, and *A*_b_. We use these values to obtain *R*_0_ = *R*_f_ + *R*_b_ + *A*_f_ + *A*_b_:

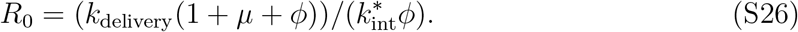

This allows us to rewrite S22-S25 as:

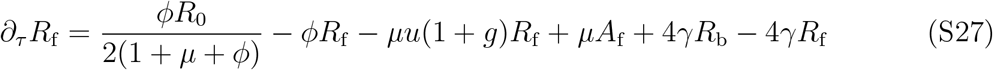

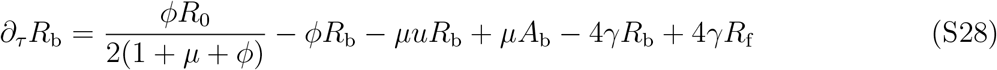

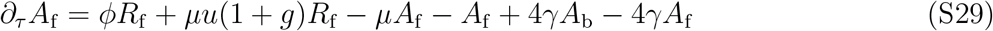

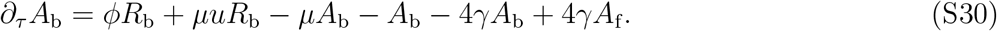

Next, we define the set point as the average activity across the two compartments:

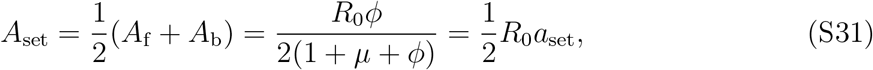

where *a*_set_ is given by equation S9. We then define the ARP as:

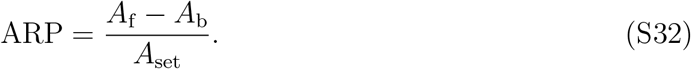

Notably, this simple model recapitulates with very high accuracy most predictions of the full model (Figure S2). Having validated the deterministic predictions of the two compartment model, we use it to explore the role of stochasticity. We note since our two compartment model describes an open system consisting of uni-molecular reactions, all species within the system will be Poisson distributed at steady state with mean values given by the deterministic equations above (76).

We construct a simple noise-aware polarization metric based on the sum of individual contributions to signaling performance from identifying either the front or the back response from the set point. To that end, we define specific stochastic realizations *κ*_f_ *∼* Poisson(*A*_f_) and *κ*_b_ *∼* Poisson(*A*_b_) of random variables describing receptor activity in the front and the back compartment respectively. We have:

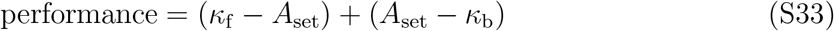

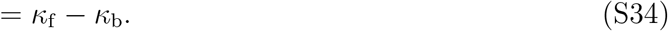

This requires *A*_f_ *≥ A*_set_ *≥ A*_b_, which is satisfied in our model (see “Analytical Model” section of Mathematica file). Failing this condition means the cell has no reference point for identifying its front or back, making the SNR zero.

Here we note *κ*_f_ and *κ*_b_ are conditionally independent Poisson variables once kinetic parameters are fixed. Their difference is described by the so-called Skellam distribution which gives the signal-to-noise relationship,

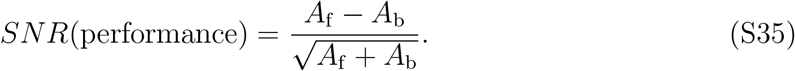

Combining equations S31 and S35 lets us rewrite *SNR*(performance) as:

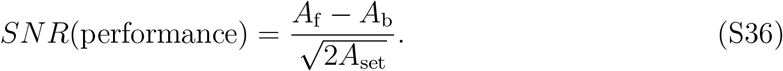

Using both equations S32 and S36 allows us to establish a simple relationship between the stochastic and deterministic metrics,

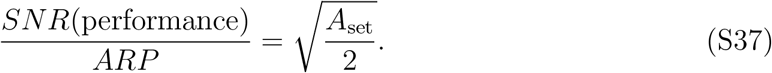

### IV. DIRECTIONAL SENSING WITH LEAKY INTEGRAL FEEDBACK

We begin with the reaction diffusion equations describing our leaky monomeric receptor system. This corresponds to a model which is nearly identical to our original model (equations S1 and S2) but with an additional reaction in which inactive receptors are internalized and degraded with a rate *k*_int_:

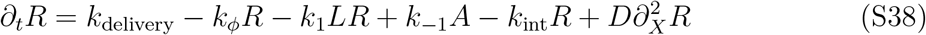

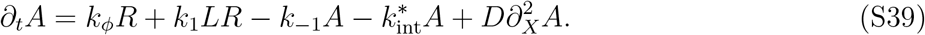

As above, in order to non-dimensionalize the model, we divide both sides of equations S38 and S39 by 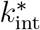, and make use of the reparameterizations: 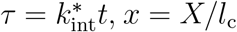 (*l*_c_ is the length of the cell), *u* = *Lk*_1_*/k*_−1_, *r* = *R/R*_0_ (*R*_0_ is the basal total receptor level), and *a* = *A/R*_0_. From this we obtain:

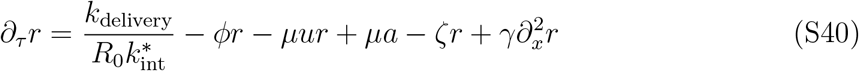

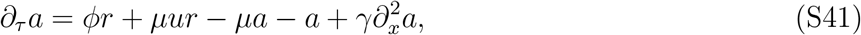

where 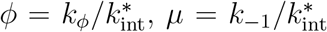, 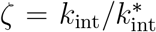, and 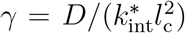. Next, in order to find *R*_0_, we solve the system described by equations S40 and S41 at steady state (*∂*_*τ*_ *r* = 0 and *∂*_*τ*_ *a* = 0) with no ligand input (*u* = 0). As above, absence of a input gradient removes any spatial variation in any receptor species along the cell, making 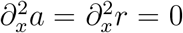. This allows this system to be solved algebraically, giving:

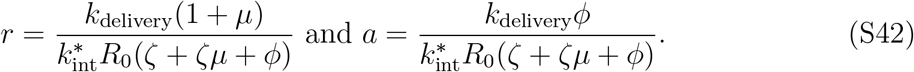

Under this dose-less steady state condition, *R*_0_ = *R* + *A* = *R*_0_(*a* + *r*), so:

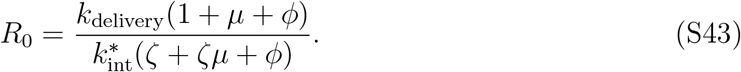

Substituting the expression for *R*_0_ back into equation S40, we obtain the following dimensionless equations for the system:

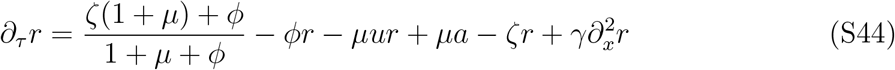

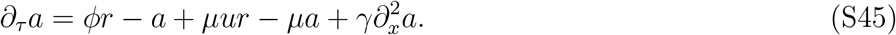

We quantify the extent of leakiness *λ* as the ratio of the internalization flux of inactive receptors compared to the internalization flux of all receptor species at steady state and in the absence of ligand input (*u* = 0):

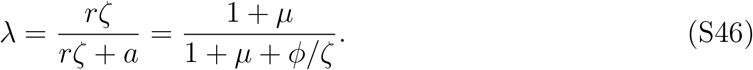

Solving equations S44 and S45 allows us to find the active receptor response profile for the leaky system. Figure S3 shows that in the absence of the leak, the back end of the cell is always below a set point (ligand-independent network activity) and the front end is always above the set point, allowing robust directional sensing over a broad range of background ligand levels. In the presence of leaky degradation, this is no longer true, i.e., the network’s activity in response to uniform ligand exposure depends on the ligand concentration and there is no true set point. Yet, we can still define directional sensing in the presence of leaky degradation by defining an effective set point (dashed black line in Figure S3), taken to be the saturated response. This set point is given by:

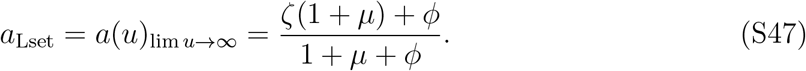

In the presence of leaky degradation, directional sensing may proceed as follows. Regions of the cell above the effective set point *a*_Lset_ are deemed to be closer to the front edge of the cell and regions below it are chosen to be closer to the trailing edge. When none of the regions on the cell are above the set point, the cell loses the ability to perform directional sensing. ARP that is cognizant of leaky degradation can be defined as follows:

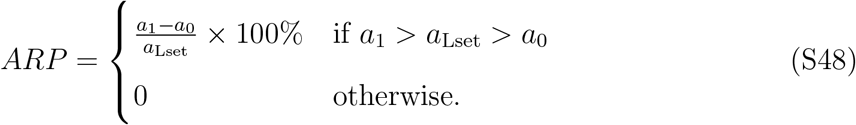

Note that this definition is inclusive of the ARP in the main text where *a*_1_ *> a*_set_ *> a*_0_ is always guaranteed (Figure S3, left panel).

Figure S4 shows the impacts of leakiness on the relative sensing range (ARP at least 80% of the maximum) and the max ARP (arrows in Figure S3). In the low leak regime, the front edge is always above the set point and the back edge is always below the set point, allowing for robust directional sensing. With an increase in leaky degradation, the front edge may start falling below the effective set point (shaded red regions in Figure S3), limiting the range of directional sensing. At very high values of the leak, no region on the cell surface will be above the set point and the cell completely loses its ability to perform directional sensing. Correspondingly, increasing leakiness decreases the range of relative sensing and correspondingly the max ARP as well. Nonetheless, negative effects of the leak are insignificant as long as the internalization flux of active receptors is kept small relative to the total receptor internalization flux (*λ ≲* 0.1) (44, 54, 72). Interestingly, increasing diffusion increases the amount of leak the system can tolerate without significant loss of performance, showing cells can limit the effects from leakiness even if they are unable to completely eliminate it.

**FIG. S2.**
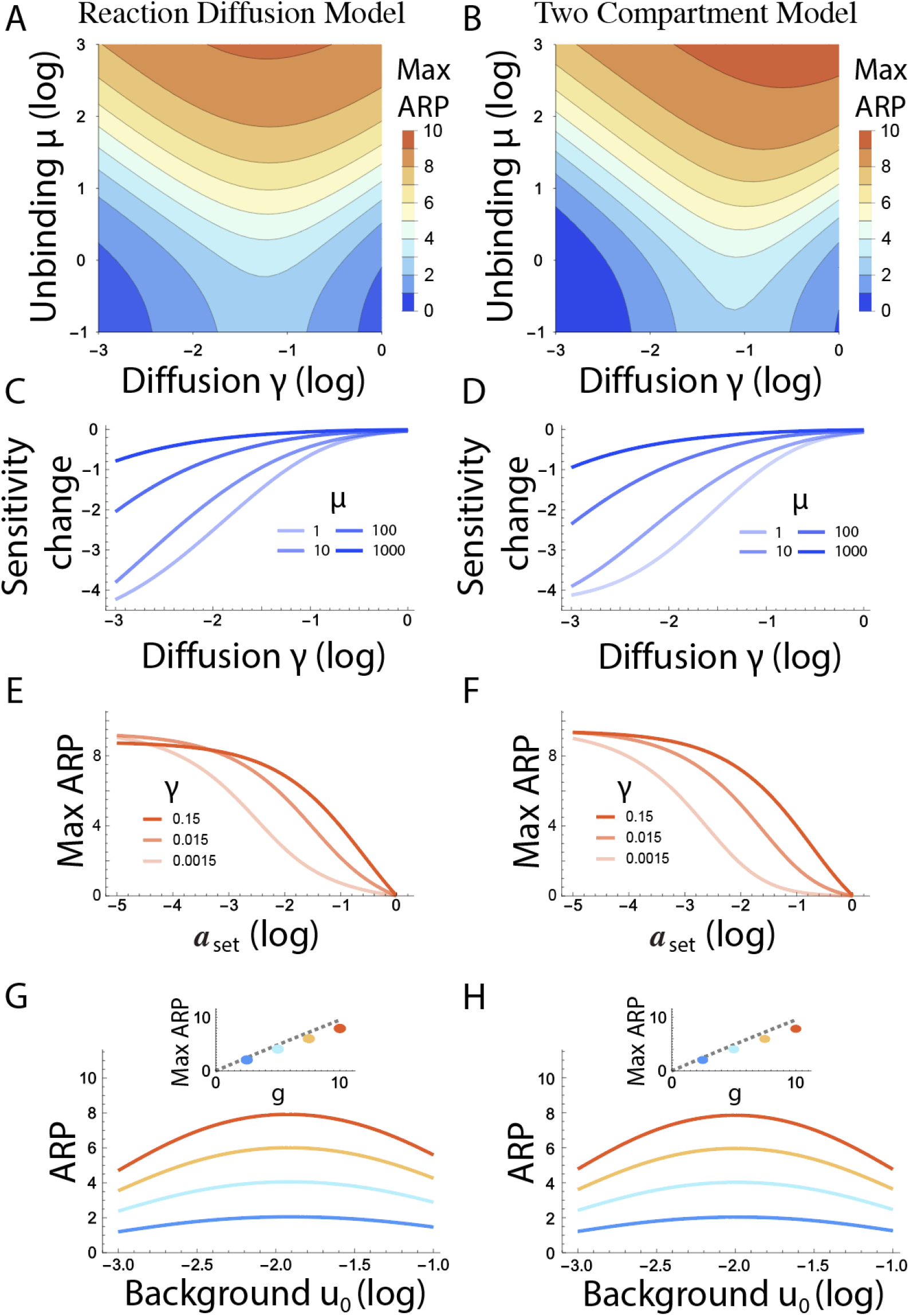
Comparing two compartment model with reaction diffusion model. (A-H) key model predictions using both the reaction diffusion model (A,C,E, and G) and the two compartment model (B,D,F, and H).

**FIG. S3.**
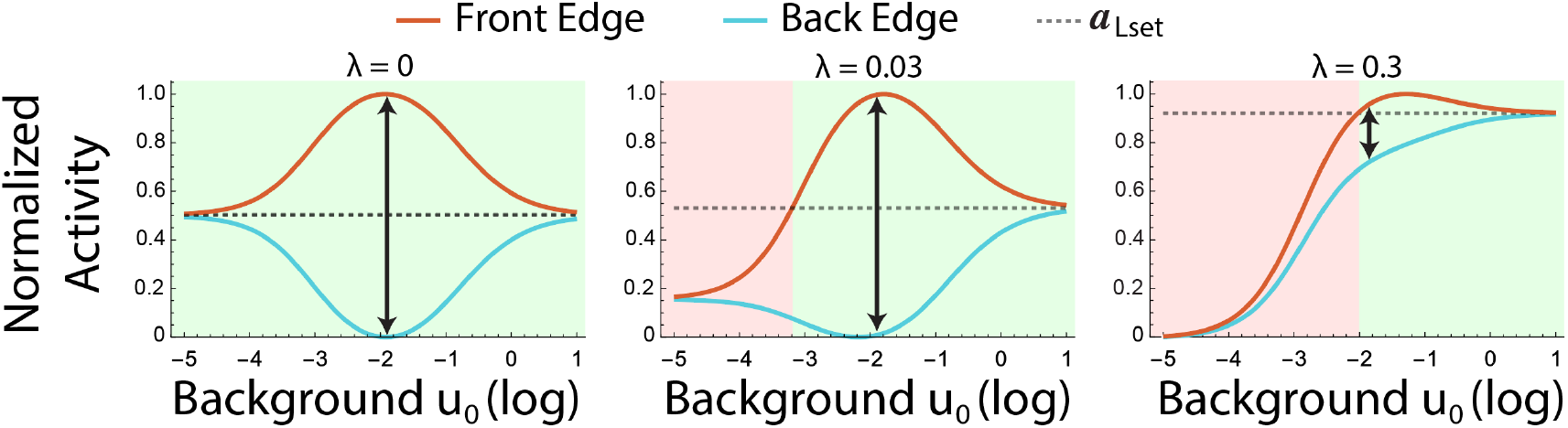
Impact of leakiness on dose response. Effects of leakiness on the responses at the front and back edge of the cell. The set point is indicated with the dashed gray line. Cells can perform robust directional sensing when the front edge is above the set point and the back edge is below it (shaded green). In contrast, the circuit is unable to detect gradients (shaded red) when the activity at both the front edge and the back edge is below this set point. The arrows show the max ARP.

**FIG. S4.**
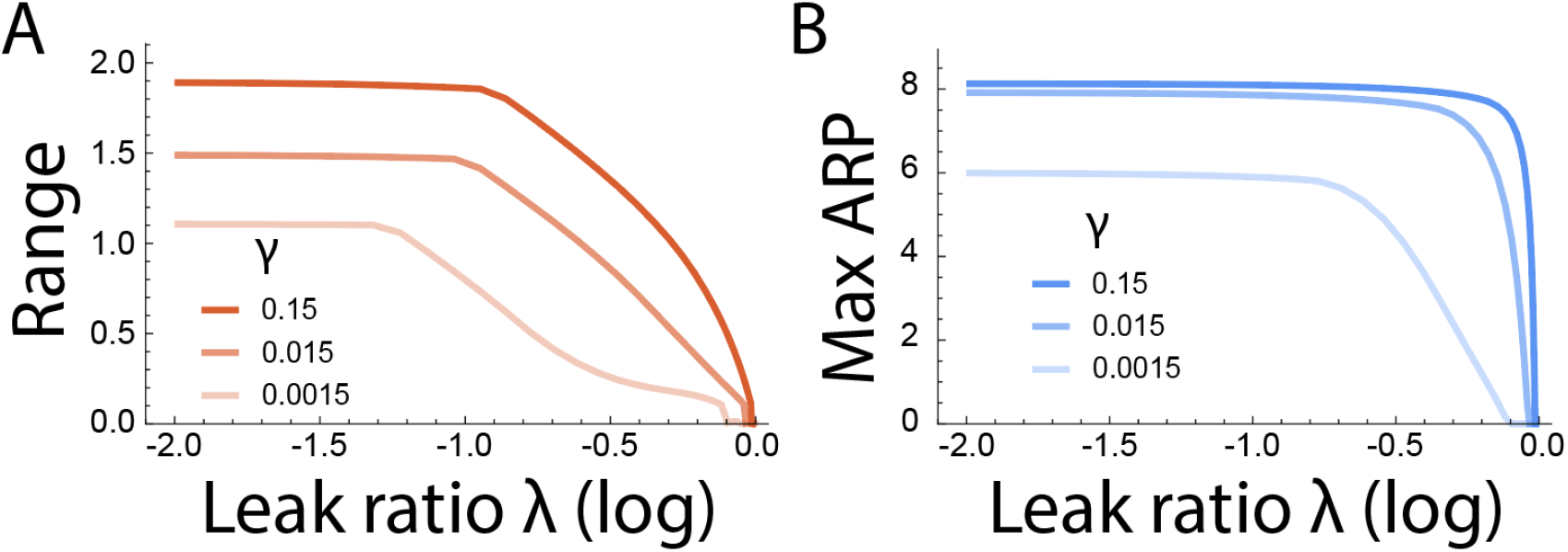
Impact of receptor leak on chemo-sensing performance. Effects of leakiness on (A) the range of relative sensing and (B) the active receptor polarization for different values of *γ*.

